# Violation of the fluctuation-response relation from a linear model of hair bundle oscillations

**DOI:** 10.1101/2022.04.15.488459

**Authors:** Florian Berger, A. J. Hudspeth

## Abstract

Spontaneous hair-bundle oscillations have been proposed to underlie the ear’s active process, which amplifies acoustic signals, sharpens frequency selectivity, and broadens the dynamic range. Although this activity is critical for proper hearing, we know very little about its energetics and its nonequilibrium properties. Systems obey fluctuation-response relations, whose violation signals nonequilibrium conditions. Here we demonstrate the violation of the fluctuation-response relation of a linear model for hair bundle oscillations. Combining analytical results with experimental data, we estimate that an energy of at least 146 *k*_B_*T* is dissipated per oscillatory cycle, implying a power output of about 5aW. Our model indicates that this dissipation attains a minimum at a certain characteristic frequency. For high frequencies, we derive a linear scaling behavior of this dissipated energy with the characteristic frequency.

## I. INTRODUCTION

One of the most fundamental properties of all living systems is their ability to use active processes to generate organization and biological function away from an equilibrium state [1]. In recent decades, tremendous advances have been made in a field called stochastic thermodynamics to describe the physics of active, fluctuating processes on a microscale [2–5]. However, applying these methods to biological systems and relating them to a biological function remains challenging.

Equilibrium systems show two important properties: they obey a time-reversal symmetry, and a fluctuation is indistinguishable from the response to a small perturbation [3, 6]. The latter property is manifested in the fluctuation-response relation that can be violated in nonequilibrium systems. Based on these two properties, different methods have been proposed to characterize and quantify active processes in biological systems [7–16].

Spontaneous oscillations of hair bundles are ideal for investigating the connection between active processes and biological function because their movement can be considered to be one-dimensional and their biological function is well-studied [17– 19]. These hair bundles protrude from the apical surfaces of hair cells in the inner ear and are the mechanosensitive elements transducing acoustic energy into electrical signals. Under appropriate ionic conditions, they display rich, non-linear spontaneous oscillations [17]. Because these oscillations violate the fluctuation-response relation, they must stem from an active process [20]. This violation of the fluctuation-response relation of oscillating bundles was restored by extending the fluctuation-response relation to nonequilibrium systems [21, 22]. Although these studies answered the question of whether these spontaneous movements were active or passive, they did not investigate the nonequilibrium energetics of the driving. Investigators have subsequently used a combination of modeling and single-cell data analysis to estimate the energy that is dissipated during an oscillatory cycle [23, 24]. These studies relied on the observed displacement of the bundle without perturbing it. As a complementary approach, the investigation of the response of a hair bundle to a perturbation potentially offers new insights into its nonequilibrium operation and provides additional information about its fascinating dynamics.

Here we analytically solve a linear model of hair bundle oscillations that adequately describes the fluctuations and responses, as observed experimentally. In such experiments, the position of the bundle is recorded and perturbed at the same time. Other degrees of freedom, for example, the cell’s ionic currents, are very difficult to access experimentally [25]. With our model, we investigate a typical experimental case, in which we can probe only the fluctuation-response relation of the bundle’s position. We derive an analytical expression for the violation of the fluctuation-response from which we determine the dissipated energy by using the Harada-Sasa equality [7]. Because this estimate for the dissipated energy is derived only from one degree of freedom of the system, it should be considered as a lower bound. After obtaining numerical values for our model’s parameters from a fit to previously published experimental data, we estimate the energy that is dissipated per cycle. The energy dissipated per cycle displays a minimum as a function of characteristic frequency. For high frequencies, we derive a linear scaling law of how the dissipated energy per cycle increases with the characteristic frequency of oscillation.

## II. VIOLATION OF THE FLUCTUATION-RESPONSE RELATION OF A SPATIAL COORDINATE

Harada and Sasa introduced an equality that relates the average energy dissipation ⟨*J*⟩ to the violation of the fluctuation-response relation of the velocity in a nonequilibrium system [7]. Considering a system without drift, the mean energy dissipation is given by

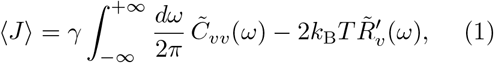

in which 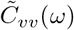 is the Fourier-transformed velocity autocorrelation function, 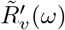 is the real part of the Fourier-transformed response function, and *γ* is a friction coefficient. This equation implies 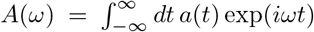 as the Fourier transformation that we will use in our study.

In contrast to the position, the velocity is not an observable that can be directly measured in experiments. Determining the velocity from a fluctuating position could lead to artifacts. To directly apply the Harada-Sasa equality to measurements of the position, we express the velocity correlations and response in terms of the position *x* correlation and response, as

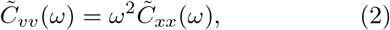

and

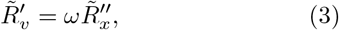

see section VII A and section VII B. Here the two primes indicate the imaginary part of the complex response function. With these transformations, we obtain the energy dissipation in terms of the correlation 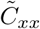 and response 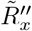 functions with respect to the position,

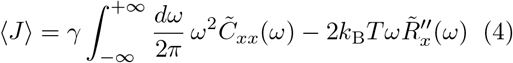

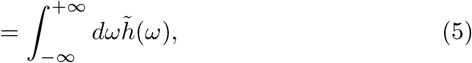

in which we introduced the violation function

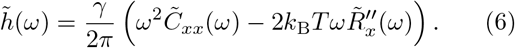

## III. VIOLATION OF THE FLUCTUATION-RESPONSE RELATION FROM A LINEAR MODEL FOR HAIR BUNDLE OSCILLATIONS

Linear models of oscillating hair bundles are sufficient to describe experimentally-observed correlation and response functions [20, 21]. Therefore, we will limit our analysis to a linear model. We describe the position *x* of a hair bundle by

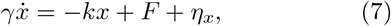

in which *γ* is an effective drag coefficient, *k* a stifness, and *F* an active driving force that is generated in the bundle. The bundle is exposed to fluctuations of the thermal environment described by the noise term *η*_*X*_. Inside the bundle, a molecular machinery generates the active force *F* that evolves in time according to

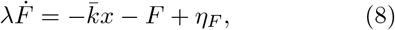

with the relaxation time *λ*, coupling constant 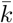, and noise *η*_*F*_ that originates from nonquilibrium fluctuations of molecular motors [26–28]. For both noise terms, we assume Gaussian noise that is delta correlated: ⟨*η*_*x*_(*t*)*η*_*x*_(0)⟩ = *D*_*x*_*δ*(*t*), ⟨*η*_*F*_ (*t*)*η*_*F*_ (0)⟩= *D*_*F*_ *δ*(*t*), with amplitude *D*_*x*_ and *D*_*F*_, respectively. Furthermore, the two noise terms are uncorrelated with each other, ⟨*η*_*x*_*η*_*F*_⟩= 0.

When we solve these equations in Fourier space for 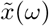, the autocorrelation is readily derived as

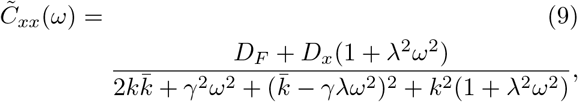

see section VII C 1. As shown in section VII C 2, we determine the linear response function of the position with the real part,

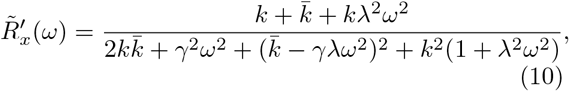

and the imaginary part,

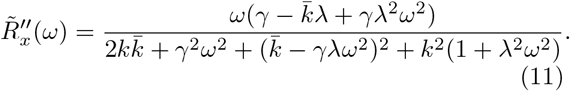

The frequency that maximizes the susceptibility 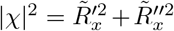 defines the characteristic frequency

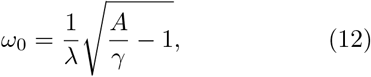

with 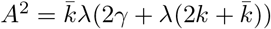.

Using eq. (11) and eq. (9), we obtain from eq. (6)the violation function

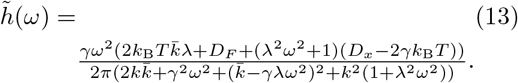

The energy dissipation as defined in eq. (4) is related to the integral 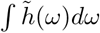 and converges to a finite value, only if 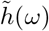 vanishes for large *ω*. Such convergence can be imposed by requiring the equilibrium fluctuation-dissipation relation for the position *x*,

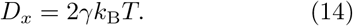

Under this condition, the violation function simplifies to

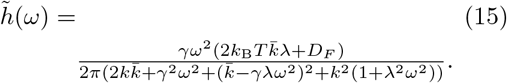

Because 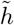 is a symmetric function, we evaluate the integral over 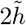 for positive values and obtain the energy dissipation

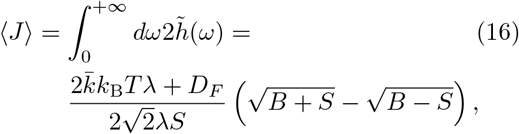

in which 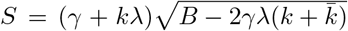 and 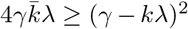, we further simplify the result to

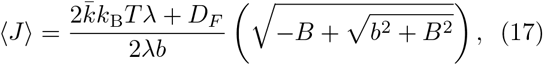

in which 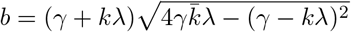.

## IV. ESTIMATING THE DISSIPATED ENERGY FROM EXPERIMENTAL DATA

To obtain numerical values for the parameter of our linear model, we fit the autocorrelation function as well as the response function to experimental data, see fig. 1(a) and fig. 1(b). The data were previously published in [20]. We fit both functions simultaneously and therefore scaled the values for each function by their respective maxima to ensure an equal weighting. From this fitting procedure, we obtained numerical values for our model parameters, given in table I. In general, the resulting fits display a good agreement with the experimental data with some deviation of the correlation function for low frequencies and of the response function for high frequencies, see fig. 1(a) and fig. 1(b). The numerical values for the parameters are similar to those obtained previously for similar model equations, but with slightly different assumptions and interpretations [20]. Although in the previous model the strength of the two noise terms was assumed to be equal, we needed to treat them independently to ensure that the integral over the violation function converged. In our case, the noise strength *D*_*x*_ of the fluctuations of the position obeys a fluctuation-dissipation relation, see eq. (14), and the source of active driving is effectively described by the noise strength *D*_*F*_ of the active force. Therefore, our numerical value for the noise strength *D*_*F*_ is about 40 times larger than in the previous study. As a further consequence, our effective coupling stiffness *k* is about five times smaller. We note that our effective model that non-reciprocally couples an active noise term to the position coordinate captures the statistics of the noise and the response very well. However, the parameters for our model should not be associated with the physical parameters that actually describe the mechanical properties of the bundle, such as the stiffness or the friction [29, 30].

**TABLE 1.** Numerical values obtained from the fits shown in fig. 1

| parameter | value from the fit |
| --- | --- |
| $\gamma$ | $8.72 \mu\text{N} \cdot \text{s} \cdot \text{m}^{-1}$ |
| $\lambda$ | $54.8 \text{ ms}$ |
| $k$ | $21 \mu\text{N} \cdot \text{m}^{-1}$ |
| $\bar{k}$ | $1.348 \text{ mN} \cdot \text{m}^{-1}$ |
| $D_F$ | $4.916 \text{ pN}^2 \cdot \text{s}$ |

**FIG. 1.**
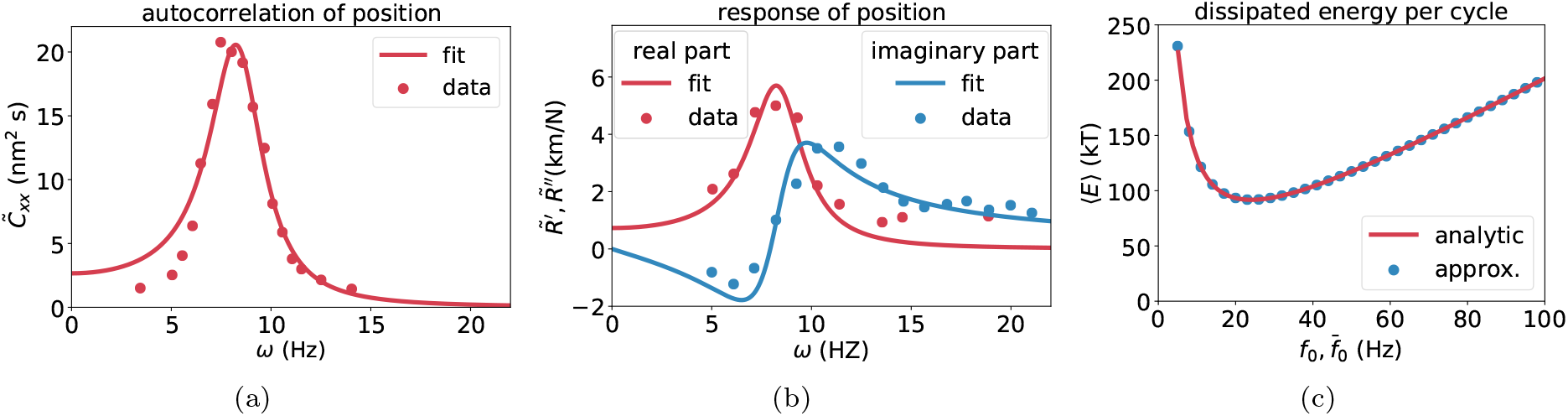
(a) The autocorrelation function of eq. (9) is fitted to experimental data previously published in [20]. The resulting numerical values for the parameters are given in table I. (b) The real (red) and imaginary parts (blue) of the response function given in eq. (10) and eq. (11) are fitted to experimental data from [20]. (c) From our model, we predict how the energy dissipation per cycle from eq. (17) behaves for different characteristic frequencies *f*_0_ = *ω*_0_*/*2*π*, with *ω*_0_ from eq. (12). We alter the characteristic frequency by changing the coupling parameter 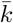 and determine the resulting parametric plot (red line). For an explicit dependency of these quantities, we derive an approximate solution of the dissipated energy per cycle in eq. (23) as a function of 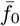 (blue dots). The energy dissipated per cycle attains a minimum and scales linearly with large characteristic frequencies of oscillation.

Using the numerical values given in table I together with *k*_B_*T* = 4.1 *·* 10^*−*21^ N *·* m in eq. (17), we obtain an estimate for the averaged dissipated energy

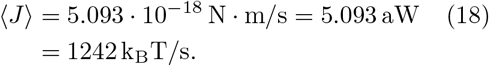

The characteristic frequency is determined from eq. (12) as

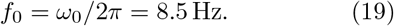

With these values, we obtain the dissipated energy per cycle as

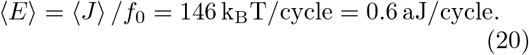

Assuming that all of this activity is related to myosin motors that hydrolysis ATP with a free energy release of roughly 10 *k*_B_*T*, we conclude that about 14 ATP molecules are hydrolyzed per oscillatory cycle [26, 27].

## V. ESTIMATED ENERGY DISSIPATION FOR DIFFERENT CHARACTERISTIC FREQUENCIES

Using our model description, we can investigate how the estimated dissipated energy per cycle depends on the characteristic frequency of the bundle. By changing the coupling strength 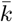 of the position coordinate to the active force, we vary the characteristic frequency *ω*_0_ of the bundle over a wide range. Considering *ω*_0_ as a variable and taking the numerical values given in table I for the other parameter values, we determine the dissipated energy per cycle as a function of the characteristic frequency in a parametric plot, see fig. 1(c). We identify a minimum of the dissipated energy per cycle at a frequency of about 20 Hz.

The numerical values for the free parameters in table I suggest that 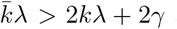 and 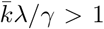. Therefore, we consider the following approximation for the characteristic frequency from eq. (12),

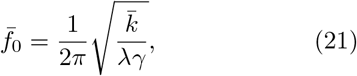

and for the energy dissipation from eq. (17),

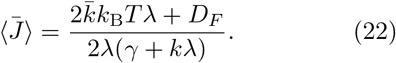

With these two equations, we express the energy dissipated per cycle as a function of the characteristic frequency,

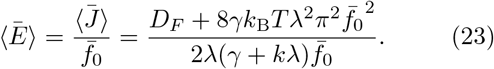

This approximation is in very good agreement with the full solution, see fig. 1(c). The minimum of this approximated dissipation function is located at the characteristic frequency

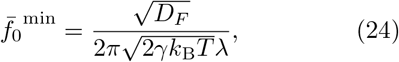

and assumes the value

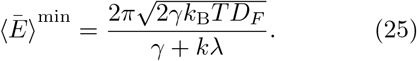

By reintroducing the noise strength *D*_*x*_ from the fluctuation-dissipation relation, we can rewrite the two equations as

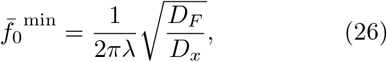

and

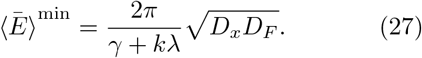

The approximation given in eq. (23) implies a linear scaling behavior for large characteristic frequencies. Our fit to the experimental data suggests that

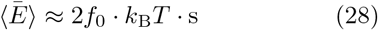

for large *f*_0_.

## VI. DISCUSSION

Spontaneous activity of hair bundles in the inner ear displays a rich variety of non-linear oscillations. Various linear and non-linear models have been introduced to describe and analyze different aspects of these oscillations [17, 28, 31–34]. The fluctuations and responses of oscillating hair bundles have been successfully described with linear models [20, 21]. Even a more complex non-linear model yielded numerically determined responses and correlation functions that were very similar [28]. This agreement suggests that a linear model is sufficient for a first analysis. In our linear model, the nonequilibrium stems from a non-reciprocal coupling between the position and an active force, whose fluctuations do not obey a fluctuation-dissipation relation. Because we determined the energy dissipation from only the violation of the fluctuation-response relation of the position, the energy that we estimated is a lower bound of the entire system. We estimated that an oscillating hair bundle generates at least 5 aW or 146 *k*_B_*T/*cycle. This value is in good agreement with a recent estimate of 100 *k*_B_*T/*cycle, based on a model fitted to bi-stable oscillations of hair bundles [24]. Furthermore, we derived a scaling behavior, indicating that the energy dissipation per cycle scaled by *k*_B ·_ *T s* is of the same order of magnitude as the characteristic frequency. Therefore, oscillations in the kilohertz range imply an energy dissipation of at least thousands of *k*_B_*T* per cycle. Manley and Gallo earlier estimated from otoacoustic emissions a power output of 141 aW per hair cell oscillating in the kilohertz regime [35]. This power output implies 34390 *k*_B_*T/*s, or tens of *k*_B_*T* per cycle, a hundred-fold difference from our estimate. Because the scaling behavior that we derived in eq. (28) relies on several assumptions from a fit to a low-frequency hair bundle oscillations, we must consider our extrapolation as a crude estimate with a large error. Nevertheless, it provides a concrete prediction of a linear scaling behavior that could, in principle, be validated experimentally by determining the fluctuation-response functions of hair cells with different characteristic frequencies. In a bullfrog’s sacculus, the frequencies of hair cells are distributed from a few hertz to 100 hertz, providing a potential system for an experimental validation of our prediction shown in fig. 1(c) [36].

We believe that a systematic study of the response functions of oscillating hair bundles will lead to a better understanding of how the active process contributes to active oscillations. It will be interesting to compare our simple description to a more refined, non-linear hair bundle models and also to experimental data. Studying response functions from hair cells with different characteristic frequencies, different oscillatory behavior, from the auditory system from different species could provide a new approach to gain insights into how our remarkable hearing is related to a nonequilibrium active process.

## ACKNOWLEDGMENTS

We would like to acknowledge fruitful discussions with Dr. U. Seifert at an early stage of this study and with Drs. R. Belousov and C. Kirst at a later stage. Possible experimental realizations were discussed with Drs. R. G. Alonso and S. Abeytunge. A.J. H. is an Investigator of Howard Hughes Medical Institute.

## VII. APPENDIX

### A. Transformation of the velocity autocorrelation into a spatial autocorrelation

Let *x*(*t*) be the position, then the spatial autocorrelation function is defined as

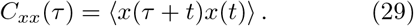

Because we assume a stationary process, the autocorrelation function is independent of *t* and depends only on the time lag *τ*. The velocity autocorrelation function is defined as

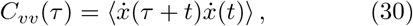

in which the dots indicate the first derivative with respect to time. We rewrite

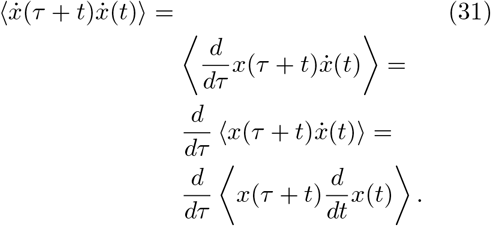

Because of stationarity, we shift the time axis to *t* = *t*^*∗*^ *− τ*, which implies a transformation of the corresponding time increments *dt* = *−dτ*, leading to

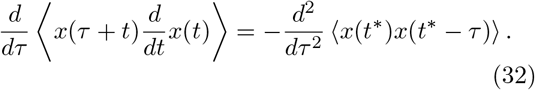

The correlation function is symmetric in time, *C*_*xx*_(*τ*) = *C*_*xx*_(*−τ*), and with *t*^*∗*^ = *t* we have

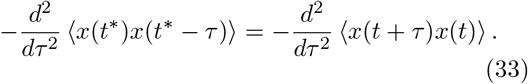

Finally, we obtain the relation

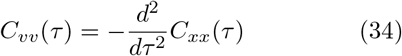

between the two correlation functions. Note that this relation can also be derived by using the Wiener-Khinchin theorem. In Fourier space, we simplify the expression to

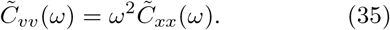

### B. Transformation of the velocity response function into a spatial response function

The linear response function *R*(*t*) of the position upon a perturbation *χ*(*t*) is defined as

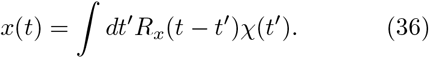

By differentiating with respect to time *t*, we obtain

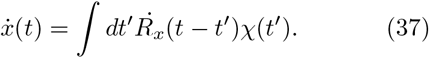

This equation defines the velocity response function

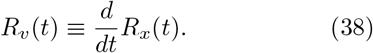

In Fourier space, this equation simplifies to

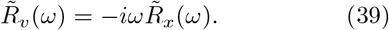

Accordingly, the real part of the velocity response reads

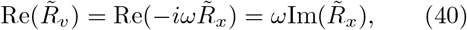

and we obtain the relation

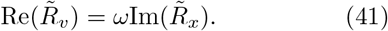

Using a prime for the real part and a double prime for the imaginary part, we obtain

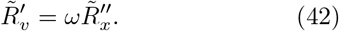

### C. Solutions of the linear model of hair bundle oscillations

We transform the model equations eq. (7) and eq. (8) to Fourier-space,

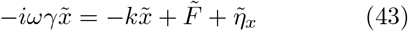

and

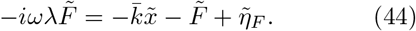

Now, we can solve for the Fourier-transformed position

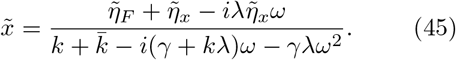

## 1. Derivation of the autocorrelation function

The averaged autocorrelation function 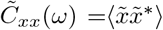 is given by the averaged product of 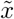 from eq. (45) with its complex-conjugate 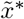

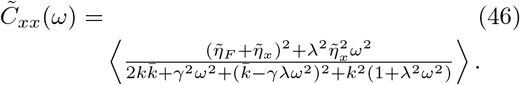

With our assumptions of the noise strengths, ⟨*η*_*x*_(*t*)*η*_*x*_(0)⟩ = ⟨*D*_*x*_*δ*(*t*), *η*_*F*_ (*t*)*η*_*F*_ (0)⟩ = *D*_*F*_ *δ*(*t*), and ⟨*η*_*x*_*η*_*F*_⟩ = 0, we derive the autocorrelation function of the position

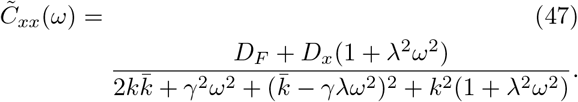

## 2. Derivation of the response function with real part

To obtain the response function, we perturb the position coordinate with a *δ*-function in the time domain, resulting in the following equations in Fourier space,

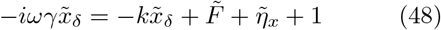

and

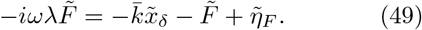

We solve for the averaged position

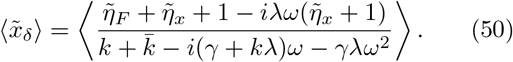

After evaluating the averages, we derive the averaged complex response

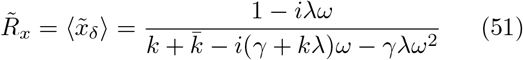

with real part

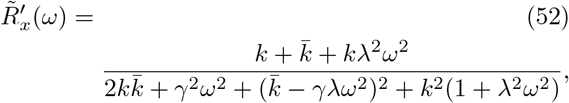

and the imaginary part

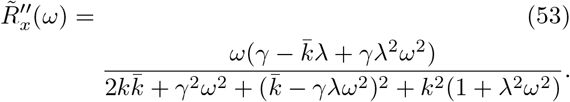

## Notes

### Competing Interest Statement

The authors have declared no competing interest.

